# Measurement and models accounting for cell death capture hidden variation in compound response

**DOI:** 10.1101/863597

**Authors:** Song Yi Bae, Ning Guan, Rui Yan, Katrina Warner, Aaron S Meyer

## Abstract

Cancer cell sensitivity or resistance is almost universally quantified through a direct or surrogate measure of cell number. However, compound responses can occur through many distinct phenotypic outcomes including changes in cell growth, apoptosis, and non-apoptotic cell death. These outcomes have distinct effects on the tumor microenvironment, immune responses, and resistance mechanisms. Here, we show that quantifying cell viability alone is insufficient to distinguish between these compound responses. Using an alternative assay and drug response analysis amenable to high-throughput measurement, we find that compounds with identical viability outcomes can have very different effects on cell growth and death. Moreover, compound pairs with additive cell growth and death effects can appear synergistic when only assessed by viability. Overall, these results demonstrate an approach to incorporating measurements of cell death when characterizing a pharmacologic response.

**Summary Points:**

- Measurements of solely live cell numbers mask important differences in compound effects.
- Additive effects on growth and death rates can appear synergistic when analyzed solely via live cell number.
- Automated imaging can provide reasonable throughput to analyze cell response in terms of cell growth and death, and endpoint analysis is similarly informative.

## Introduction

Quantifying cellular response to therapeutic compounds is essential to understanding their mechanisms of action and assessing therapeutic efficacy [1,2,3]. In the case of cancer treatments, and often with other diseases, drug activities are evaluated by quantifying the number of live cells after a short period using direct or surrogate measurements [4,5]. However, quantities beyond the number and viability of cells provide valuable information about the cellular response. Along with altering cell proliferation, promoting cell death is another important index of drug efficacy [6,7]. Incomplete eradication of drug-susceptible malignant cells allows the survival of drug-tolerant persister cell populations that can develop resistance by multiple routes [8,9,10]. Moreover, cell death can occur via a variety of mechanisms, including apoptosis and necroptosis, and selection among these outcomes can potently modulate cancer immunogenicity [11]. Limited understanding of these underlying cellular responses further complicates the assessment of drug combinations. Combination treatments are typically evaluated for their ability to enact greater effects than either compound alone [12], but typically only by quantifying viability.

Here, we show that directly measuring both cell growth and death can provide valuable information for interpreting the response of cells to single and combination treatments. We propose a framework for quantifying drug response that accounts for the compound-induced changes in rates of cell growth and death. This approach reveals extensive differences in cell response otherwise hidden by simply quantifying cell number. Of course, trade-offs exist for the breadth versus depth of analysis that can be performed to characterize cell-compound response. We show that endpoint analysis preserves much of the distinct outcomes we observe for kinetic measurements while allowing similarly high-throughput analysis to those of live cell number surrogates. These results demonstrate the need and an approach to more precisely quantify the nature of cell-compound response and interactions.

## Results

### Viability alone is insufficient to distinguish cell growth and death effects

To test whether growth and death are confounded in live cell measurements (fig. 1a), we first explored the uncertainty in a model using only these measurements. We fit typical dose-response measurements of H1299 non-small cell lung cancer cells to the chemotherapy doxorubicin (fig. 1b) to a model incorporating both cell growth and death. We assumed no cell death in the absence of drug to show the best-case scenario of assessing drug response. This model was identifiable for the live cell number (fig. 1c), and the IC_50_ and E_max_ of compound effect on cell viability were narrowly defined as 21.4 ± 1.0 nM and 26.7 ± 3.4% (90% confidence interval, CI), respectively. In contrast, the model showed large uncertainty in inferred cellular growth or death rates (fig. 1d,e). At the maximum dose, the predicted growth rate ranged 0.33–0.71 day^-1^ and death rate 0.02–0.40 day^-1^ (90% CI). The large uncertainty in outcome was due to a strong correlation in the fit values of drug effect on the growth and death rates (fig. 1f). This shows that given only live cell number quantitation one is incapable of distinguishing between reductions in cellular growth rate and increases in cell death. The number of divisions and cell deaths can vary largely while similarly fitting live cell measurements. Moreover, the number of cell divisions and cumulative dead cells can differ drastically while resulting in the same cell viability (fig. 1g).

**Figure 1:**
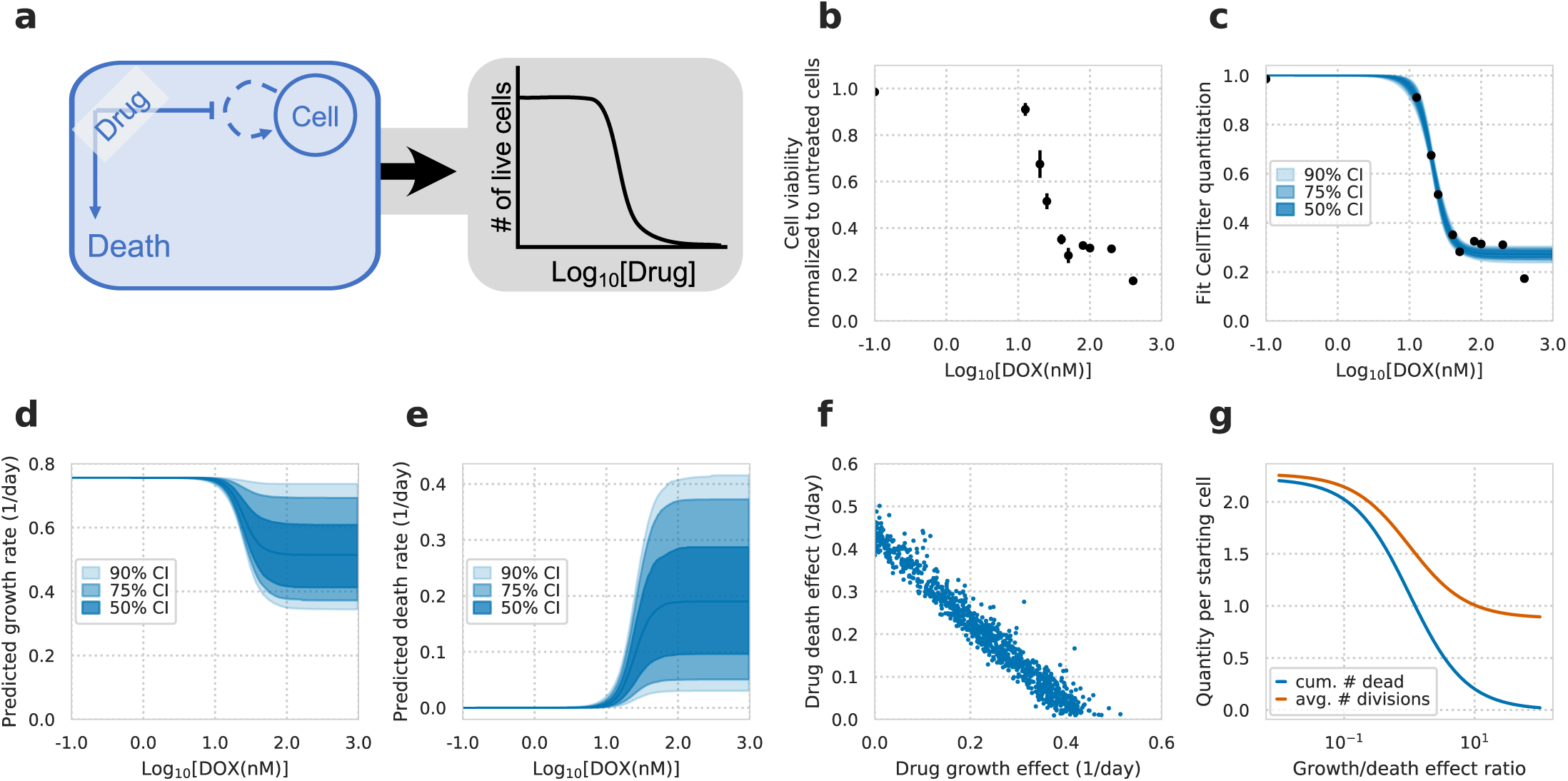
Confounding effects of cell growth and death on drug-response measurements. a, Schematic of drug response assessed by calculating relative changes in live cell number after drug treatment (gray box), and cell growth and death rate that underlie the changes (blue box). Both growth and death rates were assumed to have a Hill curve relationship to drug concentration. b, Cell viability of H1299 cells treated with doxorubicin (DOX) at varying concentrations for 72 hrs (N = 3). c, Model fit to live cell measurements. d, Model fit and confidence intervals for the predicted growth rate of cells after fitting to measurements of live cell number. e, Model fit and confidence intervals for the predicted death rate. f, Model fit posterior samples of DOX’s effect on growth versus its effect on cell death. g, Model predictions of the cumulative number of dead cells and cell divisions throughout the experiment for a constant drug effect of reducing cell viability by 75%. X-axis indicates the relative ratio in magnitude of growth versus death drug effect. Y-axis indicates the varying quantity of predicted number of cell divisions or cumulative dead cells per starting cell.

### High-throughput measurements of cell death quantify compound response

To quantify pharmacologic response, we extended our experimental measurements to those of cell death. While quantifying the number of cells over time by phase, we used an Annexin V probe to measure phosphatidylserine exposure during apoptosis and a membrane-impermeable DNA dye, YOYO-3, to measure permeabilized apoptotic and dead cells (fig. 2a and Supplementary Video 1-9) [13]. The area occupied by cells, Annexin V and YOYO-3 signal in each image were then analyzed to determine the total, apoptotic, and dead cells relative to the whole image area. We evaluated this real-time imaging method by measuring the response to doxorubicin (DOX) in H1299 cells. DOX strongly reduced the number of cells (fig. 2b, top), as seen before (fig. 1b). At the same time, we observed an increase in Annexin V while YOYO-3 increased minimally throughout the assay. Fitting these data to a model of cell growth and death (fig. 2c), we observed a strong decrease in the inferred growth rate (`div`), and only a modest increase in cell death rate (`deathRate`; fig. 2d, top). We next compared these measurements to those with another chemotherapy, vinorelbine (NVB), again observing a dose-dependent decrease in the number of live cells (fig. 2b, bottom). At the same time, we observed a large increase in both Annexin V and YOYO-3 signal. This was reflected in our subsequent analysis, inferring an increase in the death rate (fig. 2d, bottom). The fraction of cells dying through apoptosis (`apopfrac`) was also inferred to be lower in NVB as compared to DOX treatment (fig. 2d). While kinetic measurements provide a wealth of information, end-point measurement is more amenable to high-throughput experiments. We confirmed that our analysis provided qualitatively similar results using only the first and last measurements in each experiment, demonstrating that cell growth and death can both be quantified using both kinetic and endpoint measurements (fig. S1).

**Figure 2:**
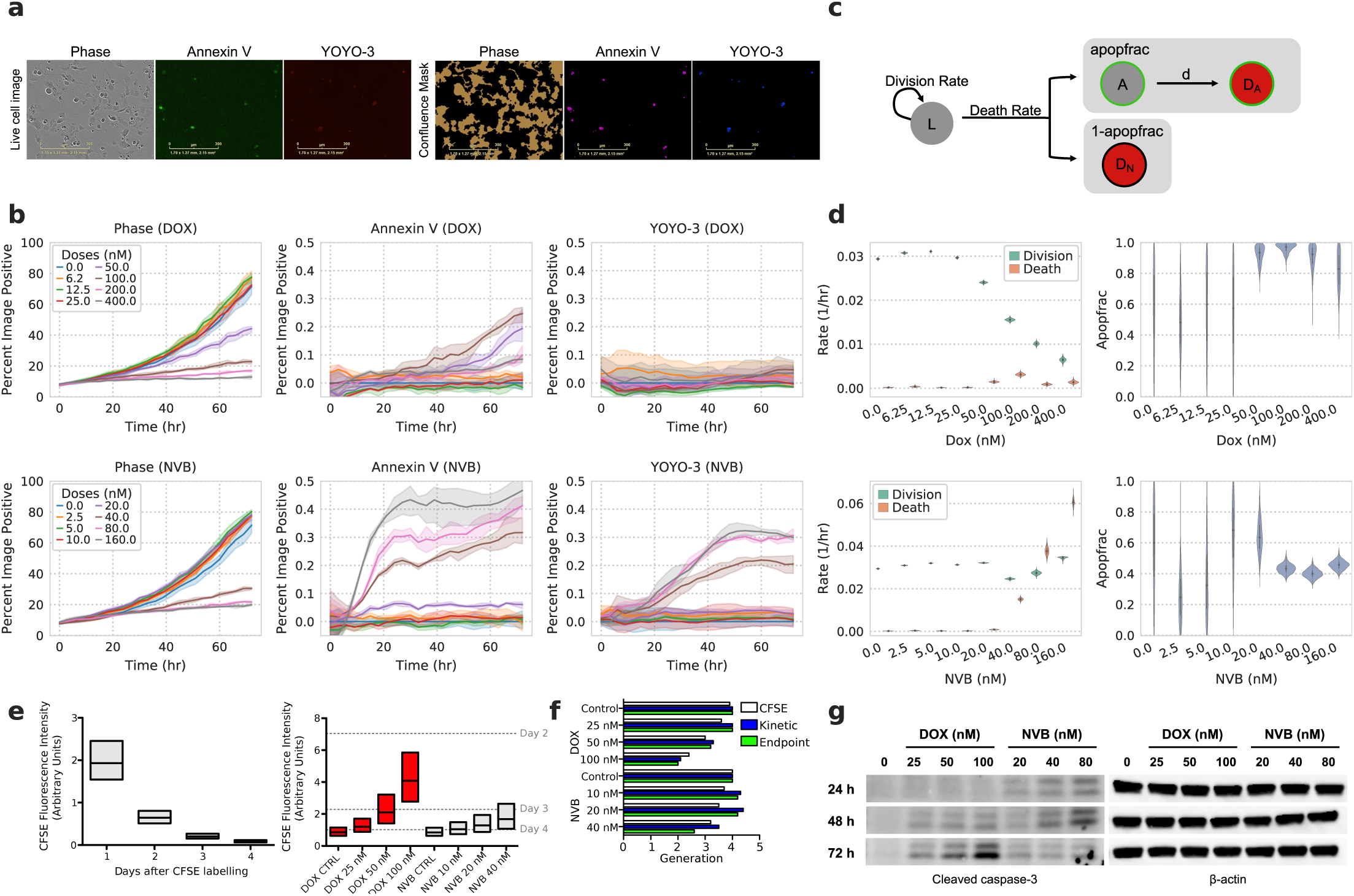
High-throughput measurements of cell death accurately quantify compound response. a, Representative images from live-cell imaging and processing. b, Experimental measurements of doxorubicin (DOX, top) and vinorelbine (NVB, bottom) response in H1299 cells over time. Phase indicates total cell confluence. Each line represents the mean of triplicate measurements for individual drug dose over time and shaded areas show the ranges of measurements. c, Schematic of the cell growth-death model. The live (`L`) cells grow at the rate of division (`div`) and die at the rate of death (`deathRate`). The model considers two fractions of cells in cell death: cells dying through apoptosis (`apopfrac`) and other modes (`1 - apopfrac`). Cells in early apoptosis (`A`) proceed into late apoptosis (`DA`) by losing membrane integrity at rate `d`. d, The model predicted `div`, `deathRate`, and `apopfrac` from the data represented in (b). Violins show posterior of model after fitting. e, Cell division analysis of CFSE-labeled H1299 cells by flow cytometry. The distribution of cells according to CFSE fluorescence intensity on indicated days for untreated cells (left) or after indicated treatments (right) for 72 hr are shown in box plots with median, lower and upper quartiles. Median CFSE intensities of 2–4 days post CFSE labeling in left graph are marked as dashed lines on the right graph. f, Number of generations for indicated conditions calculated by median fluorescence intensities from CFSE-based assay and predicted cell growth rates from kinetic and endpoint measurements. g, Western blot of cleaved-caspase 3 after treating H1299 cells with indicated drug doses for 24, 48 and 72 hrs.

To independently verify these opposing outcomes upon DOX or NVB treatment, we used a carboxyfluorescein succinimidyl ester (CFSE)-based proliferation assay to verify the distinct growth rate effects inferred by our analysis. We measured CFSE intensity of untreated cells every 24 hr starting from a day after cell labeling (fig. 2e, left). The detected intensity over time was used to estimate the number of times each cell had divided. Consistent with the inferred growth rate of our model, DOX-treated cells distributed more towards higher CFSE intensity in a dose-dependent manner, implying fewer cell divisions than with non-treated cells (fig. 2e, right). In contrast, the distribution of NVB-treated cells remained more similar to non-treated cells. Our inferred cell growth rates and CFSE measurements were overall consistent (fig. 2f).

To validate the inferred cell death rates, we measured the induction of cleaved caspase-3, an apoptotic marker, after treatment with DOX or NVB (fig. 2g). After 24-hr treatment, cleaved caspase-3 was detected in NVB-but not DOX-treated cells. Both drugs induced the caspase-3 cleavage by 48-hr treatment. These observations were consistent with the inferred cell death rates, indicating that NVB induces faster cell death than DOX. Interestingly, the level of cleaved caspase-3 after 72-hr treatment was lower in NVB-treated cells compared to DOX-treated cells. This may be explained by the lower inferred apoptosis fraction in NVB treatment while the cell death rates increased (fig. 2d) implying an increase in non-apoptotic cell death, such as caspase-independent cell death. Alternatively, a large fraction of NVB-treated cells are dead after 72 hrs. Phosphatidylserine (to which Annexin V binds) is irreversibly externalized to the cell surface by caspase-dependent scramblases during apoptosis, in contrast, it is not exposed by caspase-independent cell death [14,15], and NVB may induce caspase-independent cell death as well as apoptosis. DOX and NVB are known to operate through distinct mechanisms—inducing double-stranded breaks or preventing microtubule polymerization respectively. As a result, each compound has a differing dependency on p53 status and leads to arrest in distinct cell cycle phases, supporting that each might engage distinct cell death programs [16,17, 18]. Collectively, measuring and analyzing phase, Annexin V and YOYO-3 signals can quantify both the growth and death rate effects of drugs on cells.

### Targeted compounds also display distinct phenotypic consequences

We next evaluated whether the growth-death model can dissect the response of cancer cells to targeted compounds as well. We treated a non-small cell lung cancer cell line PC9 with 6 different targeted drugs whose effects on cell growth and death we expected to vary according to their mechanisms of action. We also used paclitaxel, a chemotherapeutic drug that is widely known to interfere with cell cycle resulting in reduced proliferation and cell death, as a standard compound for assessing the impact of targeted drugs on cell division and death rate. The total, apoptotic and dead cell measurements from each compound treatment were diverse (fig. S2) and these differences were reflected in the inferred cell division and death rates (fig. 3 and fig. S3).

**Figure 3:**
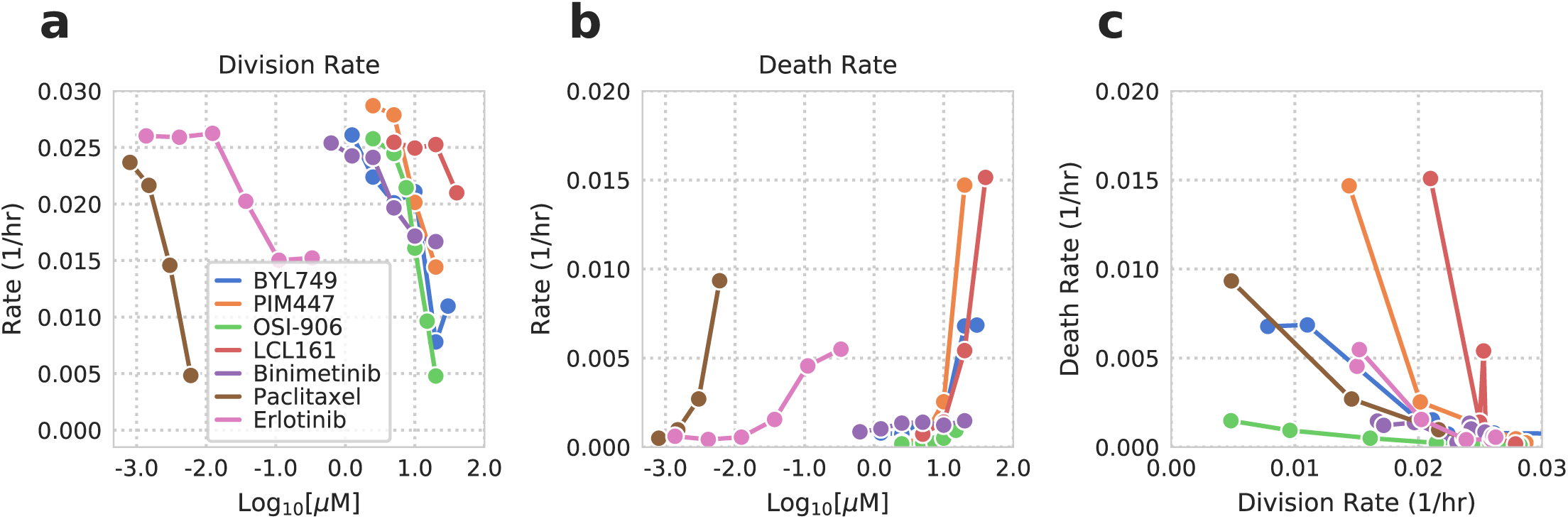
Comparing drug response for targeted compounds with distinct mechanisms of action. Model predicted cell division (a) and death rates (b) for six targeted and one chemotherapy compound. Measurements (phase, Annexin V, YOYO-3) are shown in fig. S2. The mean value of the model posterior for each dose is plotted. c, Cell division and death rates are plotted together. See fig. S3 for the probability density of each compound’s inferred `div`, `deathRate` and `apopfrac`.

We were able to classify the tested compounds into 3 types: a compound that (1) both inhibits cell division and induces cell death, (2) only inhibits cell division, and (3) only induces cell death. As expected, paclitaxel fell into the first type by simultaneously exhibiting strong cell growth suppression and death. Similar to paclitaxel, the PI3Kα inhibitor BYL749, the pan-PIM kinase inhibitor PIM447 and the EGFR tyrosine kinase inhibitor erlotinib were grouped into the first type. In contrast, both OSI-906, a dual IGF1R/INSR tyrosine kinase inhibitor, and binimetinib, a MEK1/2 inhibitor, mainly decreased cell division rates. OSI-906 inhibited the division rate as effective as paclitaxel within the range of tested doses. LCL161 is a small molecule SMAC mimetic that antagonizes multiple inhibitor of apoptosis (IAP) family proteins and augments apoptosis induction. Consistent with its mechanism of action, LCL161 showed a minimal effect on cell division while strongly enhancing cell death. Taken together, the growth-death model allows us to interpret the response of cancer cells to different targeted drugs in terms of cell division and death rates, which otherwise would not be revealed from overall phenotypic changes without a panel of experiments.

### Compounds with disparate phenotypic outcomes can appear synergistic when only analyzed by viability

Based on the changes in rate parameters by targeted drugs in fig. 3, we wondered how drugs with non-overlapping phenotypic effects might influence cell behavior when combined. We selected one compound from each of the three groups identified from fig. 3; PIM447 affected both cell division and death rates, OSI-906 affected only cell division, and LCL161 affected only cell death. Combination treatment between OSI-906 and LCL161 or PIM447 was quantified. Each compound’s effect closely matched those we expected from the single agent treatments (fig. 3, fig. 4a & f, and fig. S4a & b).

**Figure 4:**
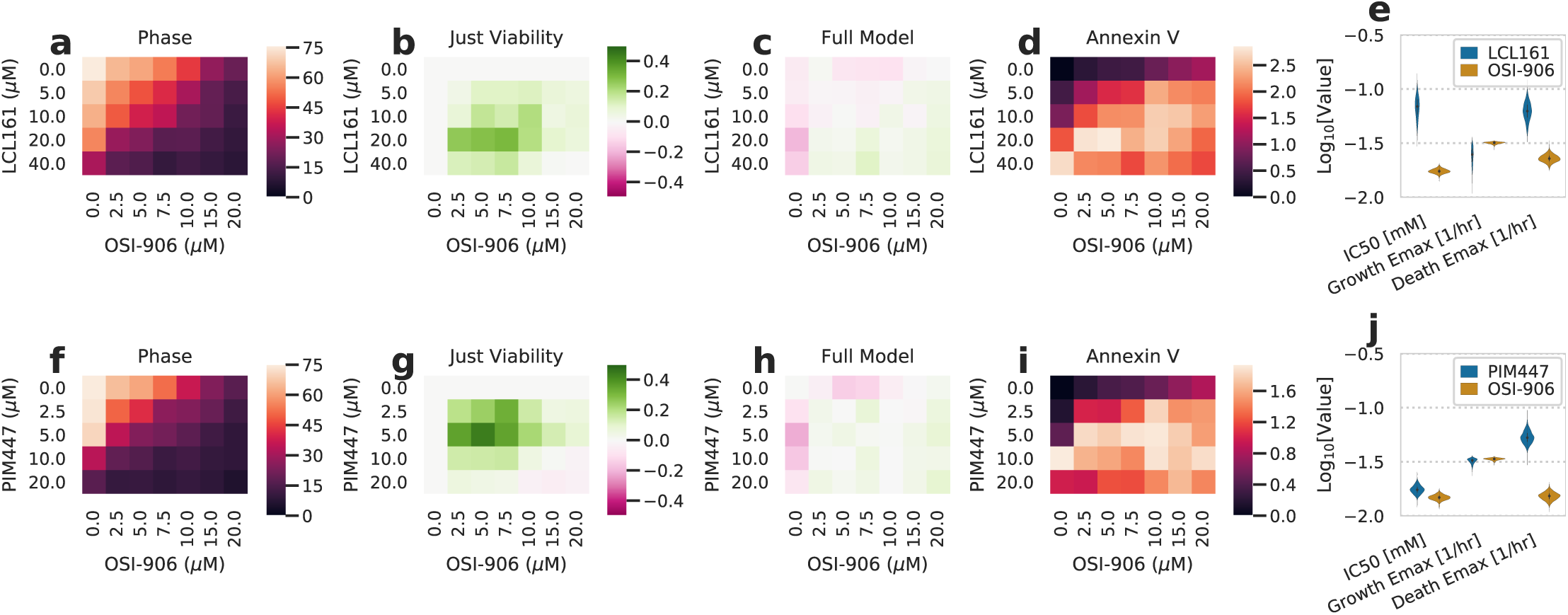
Additive interactions can appear synergistic when only assessed by viability. a & f, Heatmap of percent confluence for combinations of LCL161 (a) or PIM447 (f) with OSI-906 at 72 hrs. Color scale indicates percent confluence. b & g, Heatmap of deviation from Bliss additivity for measurements in a & f. Color scale indicates deviation from additive prediction on a scale of 0–1. c & h, Heatmap of deviation from the additive model incorporating both growth and death processes for phase measurements. Color scale indicates deviation from additive prediction on a scale of 0–1, where positive and negative values indicate drug synergy and antagonism, respectively. d & i, Heatmap of percent image area positive for Annexin V signal at 72 hrs. e & j, Fit growth rate IC_50_ and death rate EC_50_ for each drug. Fits come from the additive interaction model incorporating both processes. The experimental measurements are shown in fig. S4.

Intriguingly, the nature of each drug interaction was dependent upon whether we took cell death into account. A Bliss additivity model using just cell confluence indicated that both combinations led to a synergistic interaction (fig. 4b & g). However, we also used our combination treatment data to fit a model in which both growth and death rates respond based on a Hill curve dose-response relationship (fig. S4a & b). A model assuming Bliss additivity for the growth rate and an additive interaction for the death rate fit our measurements much more closely, despite having to account for both phase and cell death measurements (fig. 4c & h). Investigating the source of this discrepancy, we noted that the perceived synergy arose with an increase in cell death (fig. S4b/g vs. d/i). We were surprised by the difference in outcomes between each model as the IC50 and Hill coefficient of the latter model (fig. 4c & h) are assumed to be equivalent for both phenotypes. However, we expect that the difference arises from the very distinct E_max_ values for growth and death phentypes (fig. S4e & j). These terms do not simplify into one maximal effect term when entered into an exponential growth model accounting for both growth and death (methods), indicating a complex interaction between each phenotype’s effects. Therefore, we conclude that not only are measures of cell death an important component of pharmacologic response but that ignoring cell death can lead to spurious conclusions of drug synergy.

## Discussion

Here, we demonstrate how a high-throughput assay can be paired with analysis to quantify both the growth and death effects of drug response. Applying this analysis, we identify that compounds can have distinct outcomes by specifically driving growth, death, or mixed effects (fig. 3). Cell death can also take distinct forms, which can easily be quantified as relatively more or less apoptotic (fig. 2). Importantly, cell viability measurements alone cannot distinguish these divergent outcomes (fig. 1). Further, additive drug interactions in terms of cell growth or death can appear synergistic when only assessed through cell viability (fig. 4). Overall, these results show that cytotoxic drug response should be assessed by the distinct phenotypes of cell growth and death while demonstrating an approach to do so.

The approach here will benefit from improvements in the single-cell resolution of drug response and more exactly distinguishing cell death programs. Cell-to-cell variability impacts both drug response and the development of resistance [19] and single-cell technologies have enabled in-depth molecular analyses of this heterogeneity [20,21]. Within a tumor population, drug treatment can dynamically shift the balance of variability present, and so dynamic information will likely be critical [8,22]. Automated cell tracking would enable drug response quantification, with single-cell resolution, while preserving the lineage relationship of cells [23,24]. While we distinguish relatively more or less apoptotic cell death here, various cell death programs exist [25]. Improved methods to visualize distinct forms of cell death in populations of cells will allow distinct forms of cell death to be separately quantified [26].

Separating these phenotypic outcomes provides future opportunities for cancer treatment. A more detailed view of phenotypic drug response should enable treatment optimization both in a population- and patient-specific manner [27]. For maximally-effective cancer treatment, single or combination treatments should likely modulate both phenotypes. Purely cytostatic therapy leaves drug persister cells able to undergo genetic or epigenetic changes giving rise to resistance [9]. On the other hand, cell death may not overcome the replenishment of tumor cells at toxicity-limiting doses. Further, the microenvironment and host immune response to cancer are potently influenced by the routes of cell death [28,29]. These methods can be applied not only *in vitro* but in more complex, translationally-relevant models such as organoids and *in vivo* [30]. By doing so, treatments could be tailored to not just maximally reduce cell viability, but optimize cytotoxicity and cell death programs to mount a host response.

## Methods

All analysis was implemented in Python, and can be found at https://github.com/meyer-lab/ps-growth-model, release 1.0 (doi: 00.0000/arc0000000).

### Compounds and Cell culture

Doxorubicine, OSI-906, BYL719, binimetinib, and paclitaxel were purchased from LC Laboratories (Woburn, MA). PIM447 and LCL161 were obtained from Selleck Chemicals (Houston, TX). Vinorelbine was purchased from Sigma-Aldrich (St. Louis, MO). Human lung carcinoma PC9 cells were obtained from Sigma-Aldrich, and H1299 cells were provided from ATCC (Manassas, VA). All cell lines were grown in RMPI-1640 medium supplemented with 10% fetal bovine serum and 1% penicillin-streptomycin, at 37°C and 5% CO_2_.

### End-point cell viability assay and time-lapse microscopy

For the end-point cell viability assay in Fig. 1, cells were seeded at 1.5 × 10^3^ cells per well in 96-well plates and cultured overnight. Then, cells were treated with doxorubicin. After 72 hrs, CellTiter Glo reagent (Promega, Madison, WI) was added to each well and luminescence was detected according to the manufacturer’s instructions.

Cells were seeded as indicated above. The next day, each indicated treatment was added, along with IncuCyte Annexin V Green Reagent (Essen BioScience, Ann Arbor, MI) and 300 nM YOYO-3 in media containing 1.25 mM CaCl_2_. Cells were then cultured and imaged within the IncuCyte Zoom or S3 (Essen BioScience) every 3 hrs. Four fields of view were taken per well. Fluorescence images were manually thresholded and the fraction of image area with Annexin V and/or YOYO-3 signal was quantified using IncuCyte Zoom or S3 software (Essen BioScience). Finally, the fraction of area occupied by cells was analyzed by brightfield analysis.

### Hill curve identifiability model related to Fig. 1

A model of exponential growth along with death was fit to viability measurements, assuming a Hill dose-response relationship. For comparing the model to the data, the fit residuals were assumed to be normally distributed. The growth rate was measured and experimentally set to be 0.0315 1/hr, and cells were assumed to not undergo cell death in the absence of drug. The minimum growth rate (at infinite concentration of drug) was fit using a uniform prior between 0.0 and the growth rate in the absence of drug. The maximal death rate (at infinite concentration of drug) was fit using a log-normal prior of −2.0 ± 2.0 1/hr (log_10_ scale). The Hill slope was fit using a log-normal prior of 0.0 ± 1.0 (log_10_ scale). Both the IC_50_ and Hill slope were assumed to be the same for growth and death rates.

### Growth model structure

Cell behavior was modeled using a series of kinetic equations incorporating cell growth and death. We represent the overall state of a population of cells as *v* = [*L,E,D*_*a*_,*D*_*n*_], respectively indicating the number of live cells, cells within early apoptosis, dead cells via apoptosis, and dead cells via a non-apoptotic process. Using such notation, the time derivative was defined as:

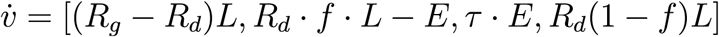

where *R*_*g*_ (or `div`) is the rate of cell division, *R*_*d*_ (or `deathRate`) is the rate of cell death, *f* (or `apopFrac`) is the fraction of dying cells which go through apoptosis, and *τ* (or `d`) determines the rate of conversion from early to late apoptosis.

If *γ* = [*R*_*g*_ − *R*_*d*_, *c* = (*R*_*d*_ · *f*)/(*g* + *d*), and *m = R*_*d*_ (1 − *f*), integrating these equations provides the solution:

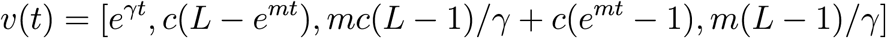

### Growth model inferrence

Predicted cell numbers were fit to experimental measurements using Markov chain Monte Carlo [31]. The percent area positive for cell confluence, Annexin V stain, or YOYO-3 stain was quantified and assumed to be proportional to the number of cells positive for each marker. Cell confluence was assumed to be the total of cells in all states. Apoptotic cells were assumed to be positive for Annexin V signal then positive for both signals after late apoptosis. Non-apoptotic cells were assumed to just be positive for YOYO-3 signal after dying. Each rate parameter was fit to the corresponding measurements within a single drug condition over time. An entire experiment, corresponding to a set of different compounds and concentrations, was fit simultaneously, allowing for a background offset and conversion factor of each quantity to be fit across the experiment.

`div` was set to have a uniform prior of 0.0–0.35 1/hr. `deathRate`, and `d` were set to have log-normal prior distributions of mean 0.001 1/hr with standard deviation 0.5 (log_10_ scale). By inspecting a calibration experiment and manually counting the cells within a field, we measured the conversion between number of cells and area of signal for the confluence, Annexin V, and YOYO-3 images. In addition, we quantified the ratio of positive area for each pair of signals when a single cell was positive for both. Each of these were set as log-normal prior distributions on the conversion values between number of cells and positive area. Finally, we observed appreciable background in the Annexin V and YOYO-3 signal, leading to signal in the absence of cells. Therefore, we set log-normal priors for the background levels with mean 0.1% of area and standard deviation of 0.1 (log_10_ scale). Each data point was assumed to have independent, normally-distributed error around the model prediction.

Sampling convergence was verified by checking that two independent runs generated insignificant differences, checking for ergodicity through the Geweke criterion comparing the first and second half of each run, and verifying an effective sample size of greater than 200. Sampling failures were solved by increasing the number of tuning samples.

### CFSE-based cell proliferation analysis

Cell division was measured using carboxyfluorescein diacetate succinimidyl ester (CFSE) dilution analysis. Cells were labeled with 5 μM CFSE (Invitrogen, Carlsbad, CA) according to the manufacturer’s protocol. The stained cells were seeded overnight in 60 mm dishes at a density of 2 × 10^5^ cells per dish, and then treated with indicated drugs next day. For 72 hrs at 24 hr intervals, cells were collected and fixed in 4% paraformaldehyde prior to acquisition on a BD LSRFortessa flow cytometer (BD Biosciences, San Jose, CA). CFSE signal intensity of 1 × 10^4^ cells was recorded and analyzed to measure cell divisions. The same cell line was labeled the day of the analysis to determine initial labeling.

### Western blot analysis

Cells were seeded at a density of 2 × 10^5^ cells per 60 mm dish 24 hrs prior to drug treatment then treated with the indicated conditions for 24, 48, and 72 hrs. After incubation, cells were lysed in 10 mM Tris-HCl pH 8.0, 1 mM EDTA, 1% Triton-X 100, 0.1% Na deoxycholate, 0.1% SDS, and 140 mM NaCl, freshly supplemented with protease and phosphatase inhibitor (Boston Bio Products, Ashland, MA). Protein concentration was measured by a bicinchoninic acid assay. 10 μg of protein from each cell lysate was subjected to SDS-PAGE, and then transferred to a polyvinylidene difluoride membrane. Each membrane was incubated overnight with antibody against cleaved caspase 3 (Cell Signaling Technology, Danvers, MA, #9664) or 1.5 hrs with HRP conjugated β-actin antibody (Cell Signaling Technology, #12262). β-actin was used as a loading control for western blot analysis.

### Drug interaction fitting

Drug interaction was assumed to follow the Bliss independence model [32]. Where indicated, this was taken to be defined as a proportional decrease in the viability of cells. That is, cell viability was normalized to 1.0 for the control condition, and then the proportional decrease in cell viability was calculated by 1.0 minus cell viability. Synergy or antagonism was identified by a greater or lesser decrease in viability than predicted, respectively.

Alternatively, Bliss additivity was defined in conjunction with a model incorporating cell death. `d` and `apopfrac` were assumed to be constant across drug concentration or combination and fit using the same prior as before. The growth rate in the absence of drug was fit using the log-normal prior of −1.5 ± 0.1/hr (log_10_ scale) based on experimental growth measurement. Cells were assumed to undergo no cell death in the absence of drug. An *E*_*max*_ of growth inhibition was fit using a Beta prior (α = 1.0, β = 1.0), where 1.0 indicates complete growth inhibition and 0.0 no growth inhibition. The *E*_*max*_ of death effect was fit using a lognormal prior of −2.0 ± 0.5/hr (log_10_ scale) where the value indicates the maximal death rate. The half-maximal concentration (*EC*_50_ or *IC*_50_) and Hill coefficient of each compound were fit using the same priors as before for these quantities and assumed to be the same for both growth and death effects.

## Acknowledgements

This work was supported by NIH U01-CA215709 to A.S.M., a Terri Brodeur Breast Cancer Foundation Fellowship to A.S.M., and in part by the UCLA Jonsson Comprehensive Cancer Center (JCCC) grant NIH P30-CA016042.

## Author contributions

A.S.M. conceived the project. A.S.M., K.W., N.G., R.Y., and S.Y.B. conducted the experiments and analyzed the results. All authors reviewed the manuscript.

## Supplementary Figures

**Figure S1:**
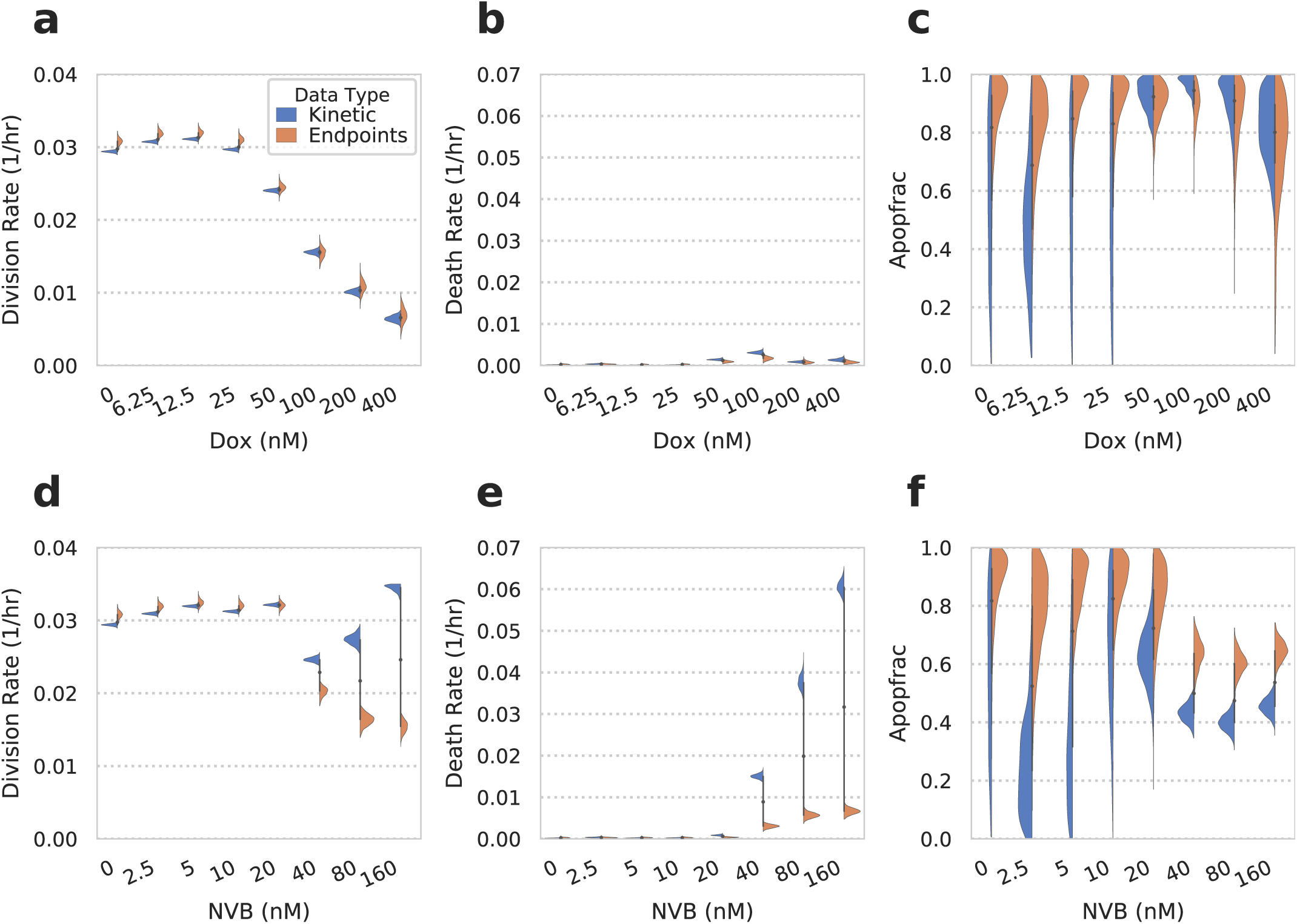
Endpoint measurements provide qualitatively similar results to those with kinetics. Violin plots indicate the model posterior after fitting. While some difference exists in the growth and death rates for high doses of NVB, the overall trends are preserved with endpoint analysis. For example, NVB more potently induces cell death than DOX, and DOX is more biased toward driving apoptotic cell death. In fact, the fits to endpoint data seem more reasonable (e.g., decreased cell division in (d)), and so the increased growth rate with kinetic measurements may be an artifact of changes in cell response over the course of the experiment.

**Figure S2:**
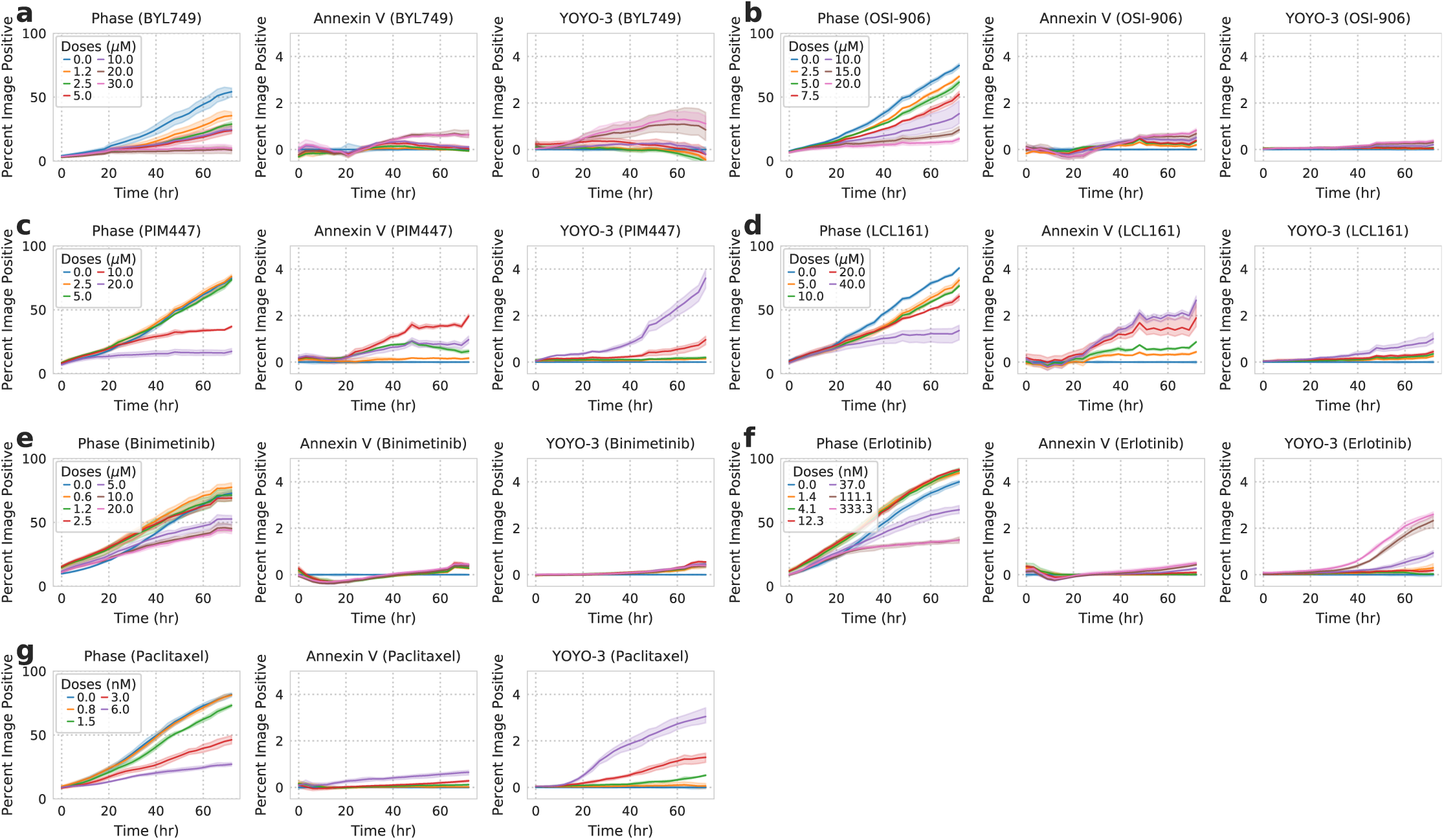
Drug response measurements for a panel of targeted agents. Model fits are shown in fig. S3, and results are summarized in fig. 3. Each line represents the mean of triplicate measurements for individual drug dose over time and shaded areas show the range of the measurements.

**Figure S3:**
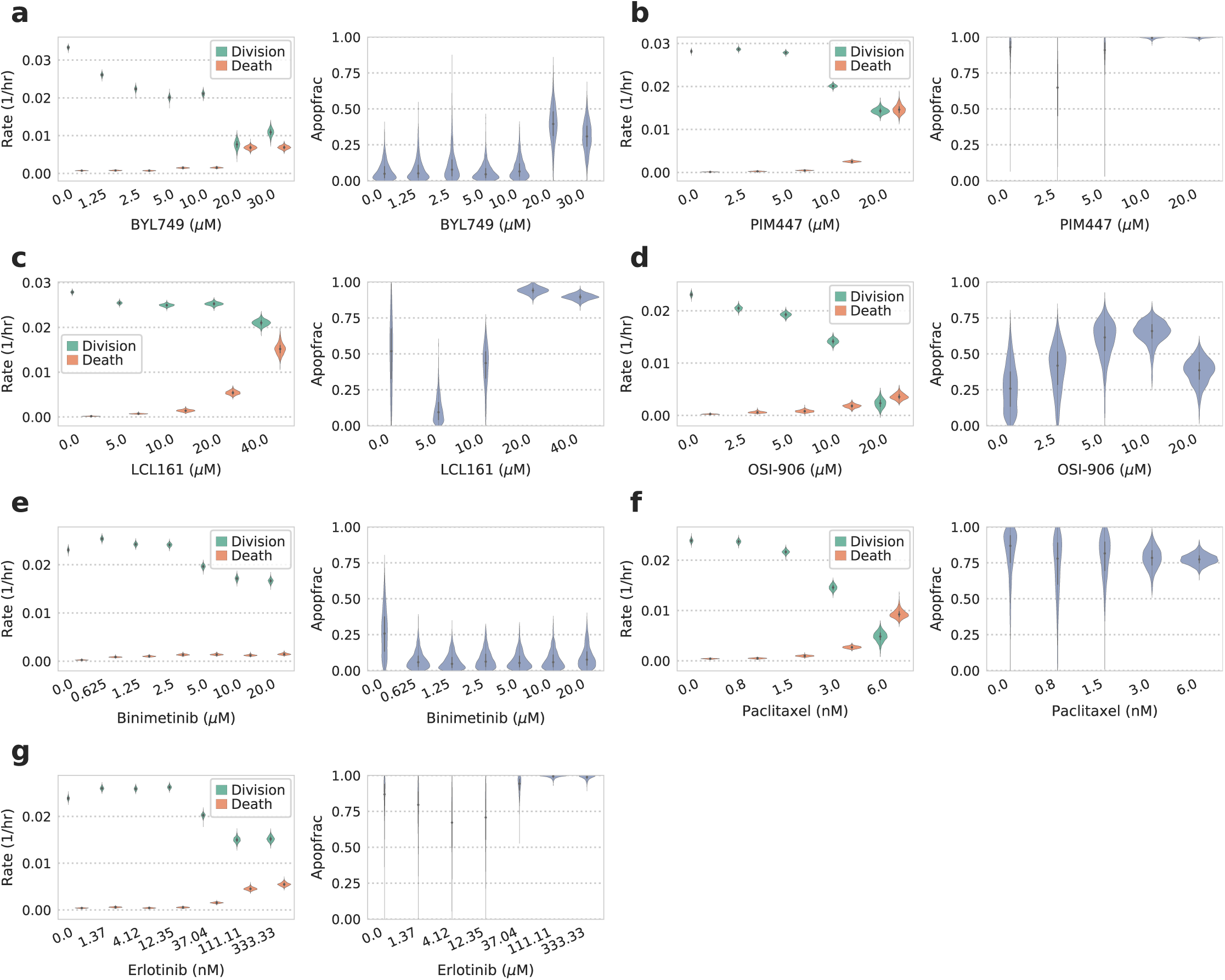
Model fitting to the drug response panel in fig. S2. Violin plots indicate the model posterior after fitting to each treatment’s kinetic measurements.

**Figure S4:**
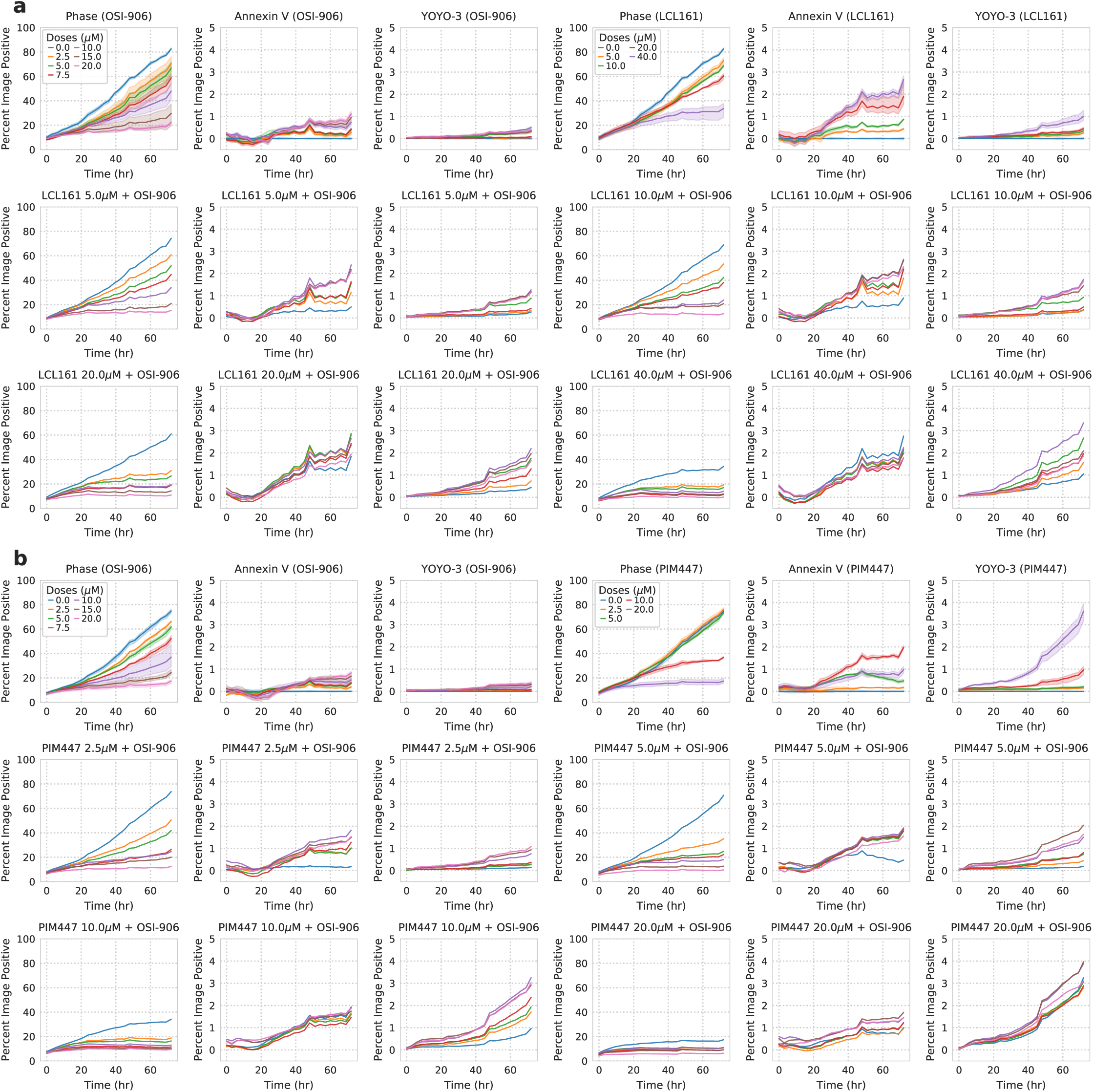
Drug response measurements for combinations of two agents with divergent phenotypic responses. Drug response measurement of LCL161 (a) or PIM447 (b) combined with OSI-906. Each line represents the mean of triplicate measurements for individual drug dose over time and shaded areas show the ranges of measurements.

